# Pathogenic transcriptional reprogramming of fibro-adipogenic progenitors in mice with FOP and its mitigation by inhibition of ACVR1 and activin A

**DOI:** 10.64898/2026.09.23.752902

**Authors:** Sean J. Stoessel, Lorraine N. Burdick, Alison J. Davis, David J. Goldhamer

## Abstract

Individuals with fibrodysplasia ossificans progressiva (FOP), a rare genetic disorder caused by mutations in the bone morphogenetic protein receptor ACVR1 (also known as ALK2), experience progressive and severely debilitating endochondral heterotopic ossification (HO). Previous studies have identified fibro-adipogenic progenitors (FAPs) as a major cellular source of HO in FOP mouse models. Here, we used mRNA sequencing of FAPs isolated from injured muscle of FOP mice to capture the earliest gene expression changes associated with the transformation of muscle tissues into bone. Whereas the transcriptomes of mutant and wild-type FAPs were indistinguishable prior to injury and at 1 day post injury, dramatic dysregulation of gene expression was observed at 3 and 5 days after injury. Surprisingly, gene expression changes suggested early specification of osteochondral lineages by 3 days post injury, approximately 3 to 4 days before the appearance of histologically identifiable cartilage. We identified dysregulation of genes encoding components of major signaling pathways, cell cycle regulators, extracellular matrix components, and hypoxia response mediators, and showed that these gene expression changes are temporally associated with abnormal population growth of mutant FAPs. We further demonstrated that treatment of FOP mice with either an anti-activin A monoclonal antibody or the ACVR1 kinase inhibitor BLU-782 (IPN60130; fidrisertib) normalized the expression of the majority of dysregulated FAP genes. However, the relative efficacy of the drugs differed, and distinct sets of overlapping genes were normalized by the treatments. The substantial transcriptional changes caused by these drugs in FAPs from normally regenerating muscle were also enumerated and described. These findings define early transcriptional events associated with pathogenic FAP reprogramming and provide a molecular framework for understanding the shared and distinct effects of activin A and ACVR1 inhibition.

**One Sentence Summary:** Early transcriptional reprogramming of fibro-adipogenic progenitors in FOP mice

## INTRODUCTION

Skeletal muscle injury elicits a series of tightly coordinated molecular and cellular events that ultimately result in muscle regeneration and the restoration of muscle structure and function. Interference with the function of muscle stem cells (satellite cells) through genetic means or alterations in the muscle signaling environment leads to failed or impaired regeneration. Among the extrinsic factors required for a normal regenerative response is a robust, but highly regulated, inflammatory response, and the action of tissue-resident stem/progenitor cells that help provide an appropriate signaling environment for normal satellite cell function and tissue homeostasis(^1–3^). In this latter regard, fibro-adipogenic progenitors (FAPs) – multipotent mesenchymal cells resident in the interstitium of skeletal muscle and associated soft tissues(^4,5^) – are positive effectors of muscle regeneration under normal physiological conditions(^4,6–10^). FAPs produce several well-established positive regulators of regeneration(^4^), including IL-6, IGF-1, WNT1, WNT3A, and WNT5A, and co-culturing experiments suggest that FAPs can support satellite cell expansion(^7^) and differentiation(^4^). FAPs also support regeneration by dynamic remodeling of the extracellular matrix (ECM) following muscle injury(^7,9^). Indeed, depletion of PDGFRα+ FAPs by diphtheria toxin-mediated ablation impairs regeneration(^11^).

Under pathological conditions in which muscle regenerative capacity is impaired, FAPs are likely the primary source of adipocyte and fibrotic accumulations, which can severely disrupt muscle architecture and function(^4–7,9,12–14^). Endochondral bone formation in skeletal muscle and associated soft tissues (heterotopic ossification: HO) is a distinct pathological outcome that can result from muscle injury, and FAPs are a source of cartilage and bone in mouse models of HO. For example, lineage-tracing studies have shown that wild-type (WT) FAPs can become programmed for skeletal differentiation following intramuscular injection of osteogenic bone morphogenetic proteins (BMPs)(^12,15,16^), or in the context of an abnormal inflammatory response in the absence of exogenous BMPs(^17^). In fact, following BMP2 injection, ∼80% of heterotopic bone derives from PDGFRα+ FAPs(^17^), arguing that FAPs may represent the major BMP-responsive skeletal progenitor resident in muscle tissue(^12,17^). FAPs are also a major source of skeletal progenitors in mouse models of the rare autosomal-dominant genetic disorder fibrodysplasia ossificans progresssiva (FOP), which is characterized by aggressive and progressive HO of skeletal muscles, tendons, and ligaments. The cumulative effects of HO, which typically begins in early childhood(^18^), include ankylosis of major joints, muscle loss, and other complications that cause pain, severely restricted mobility, and reduced life expectancy(^19–21^). The great majority of cases of FOP with classical clinical features result from a single arginine to histidine (R206H) substitution in the highly conserved glycine-serine domain of the BMP type I receptor, ACVR1(^19–21^). Mice engineered to express ACVR1(R206H) in FAPs (R206H-FAPs) using Tie2-Cre(^22^) or Pdgfra-Cre(^23^) drivers develop HO after muscle injury (both Cre drivers)(^24–26^), or exhibit juvenile-onset spontaneous HO (Pdgfra-Cre)(^24,27^), collectively recapitulating most major manifestations of the disease. In contrast to the extrinsic effects of the muscle environment in controlling FAP differentiation noted above, ACVR1(R206H) can co-opt FAPs for skeletogenic differentiation by cell-autonomous mechanisms(^24^). Notably, lineage-tracing studies have shown that Tie2+ embryonic progenitors do not significantly contribute to development of the normal skeleton(^12,15,24,27^). Thus, reprogramming of FAPs for pathological skeletogenesis may utilize novel genes and pathways, particularly upstream of expression of canonical drivers of cartilage and bone differentiation and maturation.

While ACVR1(R206H) remains responsive to BMP ligands(^24–26,28,29^), this single amino acid change alters signaling specificity such that an osteogenic response is now elicited by activin ligands(^28,29^). Importantly, systemic delivery of a blocking antibody to activin A inhibits both injury-induced and spontaneous HO, demonstrating that activin A is an essential ligand for HO in FOP mice(^24,25,27,29^). Highly selective small-molecule inhibitors of ACVR1 also inhibit HO in FOP mice(^30^), and both strategies are being evaluated in clinical trials. In the present study, we used bulk mRNA sequencing (RNA-seq) to define the time course of transcriptional reprogramming of R206H-FAPs and determine the degree to which blocking activin A or inhibition of ACVR1(R206H) kinase activity normalizes the R206H-FAP transcriptome and population growth during early post-injury stages. The effects of drug treatment on FAP gene expression during muscle regeneration in WT mice were also defined. These data identify presumptively dysregulated biological processes and pathways that likely contribute to the cellular and tissue-level pathological response to muscle injury in FOP mice and provide comprehensive and sensitive molecular metrics of drug efficacy for these therapeutic modalities.

## MATERIALS AND METHODS

### Mice and genotypes

All animal procedures were approved by the University of Connecticut’s Institutional Animal Care and Use Committee (IACUC). 2-3-month-old female FOP (*Acvr1^tnR206H/+^*;*R26^NG/+^*; Tie2-Cre) and control (*R26^NG/+^*;Tie2-Cre) mice were used for these studies. The exclusive use of females was based on recent findings that female FOP mice produce greater average HO volumes following muscle injury than male FOP mice(^31^). All mice were on a highly enriched FVB background.

### Skeletal muscle pinch injury

Prior to pinch injury, mice were sedated with isoflurane and the lower hindlimbs were shaved. To induce a reproducible muscle injury, a modified Randall Selitto Paw Pressure Test Apparatus (IITC Life Science)(^24^) was used to apply roughly 4,000 grams of force to both sides of the posterior muscle compartment of the lower hindlimbs (two consecutive pinches for 5 sec each).

### Drug dosing regimens and endpoints

A working solution of BLU-782, supplied by Blueprint Medicines Corporation, was made weekly by dissolving the powder form in 20% (2-hydroxypropyl)-β-cyclodextrin in citrate buffer at pH 4.5 to a concentration of 5 mg/mL and aliquoted into single-use tubes, as instructed by Blueprint Medicines. Beginning 3 days prior to muscle injury and extending until 1 day prior to tissue collection, mice received daily doses of 50 mg/kg BLU-782 via oral gavage. Untreated FOP mice were not dosed with vehicle to avoid the possibility of inducing even mild injury through handling. Mice treated with ActA-mAb (1 mg/mL in sterile PBS; supplied by Acceleron Pharma) received a single subcutaneous injection at a dose of 10 mg/kg at the time of muscle injury. These dosing methods are highly effective at inhibiting HO(^24,30^).

Untreated and treated FOP mice were collected at 0, 1, 3, and 5 days post injury (dpi). Untreated control mice were collected at the same post-injury time points, whereas control mice treated with ActA-mAb or BLU-782 were collected only at 5 dpi. Biological triplicates were used for each comparison group. Muscle from two mice was pooled for each biological replicate of the uninjured and 1 dpi groups, as FAP yields were insufficient from one mouse for accurate RNA quantification and subsequent cDNA library preparation.

### µCT imaging

µCT imaging was performed using the IVIS SpectrumCT (PerkinElmer) at medium resolution, which acquires 75 µm voxel size images using estimated radiation doses of 132 mGy for a 3.5 min scan time. Mice were anesthetized with isoflurane during imaging.

### Tissue dissociation and flow cytometry/fluorescence-activated cell sorting

For FAP quantification and collection of FAPs at 3 and 5 dpi for bulk mRNA-seq, muscle tissue was minced for 4-5 min and then enzymatically digested with 700 U/mL collagenase II and 0.3 mg/mL dispase in Dulbecco’s Modified Eagle Medium (DMEM) for 1 hr at 37° C. Tissues were triturated for 30 sec every 15 min during incubation. Cell suspensions were passed through a 70 µm filter and centrifuged at 800 x g for 5 min at 4° C. Cell pellets were washed twice with cold DPBS before antibody staining. The following modifications were employed to improve FAP yields at 0 and 1 dpi. Muscle was gently teased apart along the axis of myofibers for 1 min instead of mincing the tissue. Rather than dissociating the muscle by trituration, the tissue was slowly rotated in an incubator at 37° C for 45 min. The relative centrifugal force was reduced to 305 x g, and 50 µg/mL of DNase I was added to the final cell suspension to prevent extracellular DNA from clumping cells or non-specifically binding antibody.

Washed cell suspensions were incubated with anti-CD140a (PDGFRA)-APC (eBioscience, 17-1401-81; 1:100), anti-Ly6A/E (SCA-1)-v450 (BD Biosciences, 560653; 1:400), anti-CD31-bv711 (BD Biosciences, 740690; 1:800), and anti-CD45-bv711 (BD Biosciences, 563709; 1:500) antibodies in 2% fetal bovine serum in DPBS for 30 min on ice. Sorting was performed on a FACSAria II (BD Biosciences) equipped with 405, 488, 561, and 640 nm lasers using a gating strategy that was previously described(^25^). An LSRFortessa X-20 (BD Biosciences) was used for flow cytometry analysis using the same gating strategy and laser wavelengths. Precision count beads (BioLegend) were added at known concentrations (verified using a hemacytometer) to allow calculation of the total number of a target cell population within the sample.

### RNA isolation, cDNA library preparation, and bulk mRNA-sequencing

FAPs were sorted directly into RLT lysis buffer (Qiagen), vortexed for 30 sec, and kept on ice. The RNeasy mini kit protocol was used to column-capture total RNA. RNA quality was confirmed (RIN ≥ 8.0) and quantified using an Agilent Tapestation 4200 with high-sensitivity RNA screen tape. TruSeq Stranded mRNA library preparation (Illumina) was used to capture mRNA and create cDNA libraries, which were then sequenced using either the NextSeq 550 or NovaSeq 6000 (Illumina). Approximately 30 million 75 bp, single-end reads were generated per sample.

### Differential gene expression and gene ontology analyses

The computational workflow consisted of read quality control using FastQC and MultiQC, read quality filtering using fastx-toolkit, alignment to an annotated mouse reference genome (Mus_musculus.GRCm39.105) using the splice-aware aligner, Hisat2, generation of a sorted bam file using SAMtools, and count matrix generation using featureCounts(^32^). Qualimap and SAMtools-stats were used to assess mapping efficiency, multimapping, and 5’-3’ bias of reads(^33,34^).

DESeq2 was used to perform independent gene filtering, dispersion estimates, normalization, and statistical testing for differentially expressed gene analysis(^35^). PANTHER was used to perform statistical overrepresentation analysis on differentially expressed genes(^36^). Ingenuity Pathway Analysis (IPA) (Qiagen) and ShinyGO v0.80(^37^) was used to confirm PANTHER results. All heatmaps, bar plots, PCA plots, and MA plots were created in R or Graphpad Prism.

### Statistical analysis

Differences in calculated FAP numbers from flow cytometry were tested for significance using a one-way ANOVA with Tukey’s multiple testing correction. For RNA-seq differential gene expression analysis, DESeq2 was used for pairwise comparisons. The Wald test followed by the Benjamini-Hochberg correction for multiple testing was used to determine significance at an adjusted p-value of < 0.05 using DESeq2 normalized read counts. To be considered differentially expressed, gene expression changes required log2 fold changes in mean normalized counts of ± 0.585 (at least 50% higher or 33% lower than the reference).

## RESULTS

### A small-molecule kinase inhibitor of ACVR1 is highly effective at blocking FAP-mediated HO in FOP mice

Activin A blocking antibodies (ActA-mAb) are highly effective at inhibiting both spontaneous and injury-induced HO in FOP mouse models(^24,29^). We sought to determine whether the highly specific small-molecule kinase inhibitor of ACVR1, BLU-782 (developed by Blueprint Medicines), which was shown to be effective at blocking HO in an independently-derived FOP mouse model(^30^), was similarly effective at blocking HO in our injury-inducible FOP model (*Acvr1^tnR206H/+^*;Tie2-Cre). Female FOP mice between 2 and 3 months of age were pinch-injured as previously described(^24^), and 50 mg/kg BLU-782 was administered daily via oral gavage, beginning 3 days prior to injury (n = 6). In a previous study, daily dosing with 50 mg/kg BLU-782 was shown to be 100% efficacious in blocking HO(^30^). Only female mice were used for this study, as females display a significantly greater HO response at endpoint in this model(^31^). As assessed by µCT imaging at 14 dpi, limbs of untreated FOP mice were heavily burdened with heterotopic bone (Fig. 1A, n = 5), whereas BLU-782 treatment resulted in near-complete inhibition of HO (Fig. 1B, n = 6). A minority of injured limbs of both BLU-782-treated and untreated FOP mice exhibited minor exostosis of the distal tibia (Fig. S1), bone overgrowth that may have resulted from tensile forces indirectly applied to the periosteum or associated connective tissues during the muscle pinch-injury procedure. Whether the inability of BLU-782 to block exostosis reflects limited drug bioavailability or the engagement of additional signaling pathways requires clarification. We note, however, that FOP mice subjected to cardiotoxin-induced muscle injury also develop exostosis whereas control mice lacking ACVR1(R206H) do not(^26^), indicating that ACVR1(R206H) activity is required for development of exostosis in these muscle injury models.

**Figure 1.**
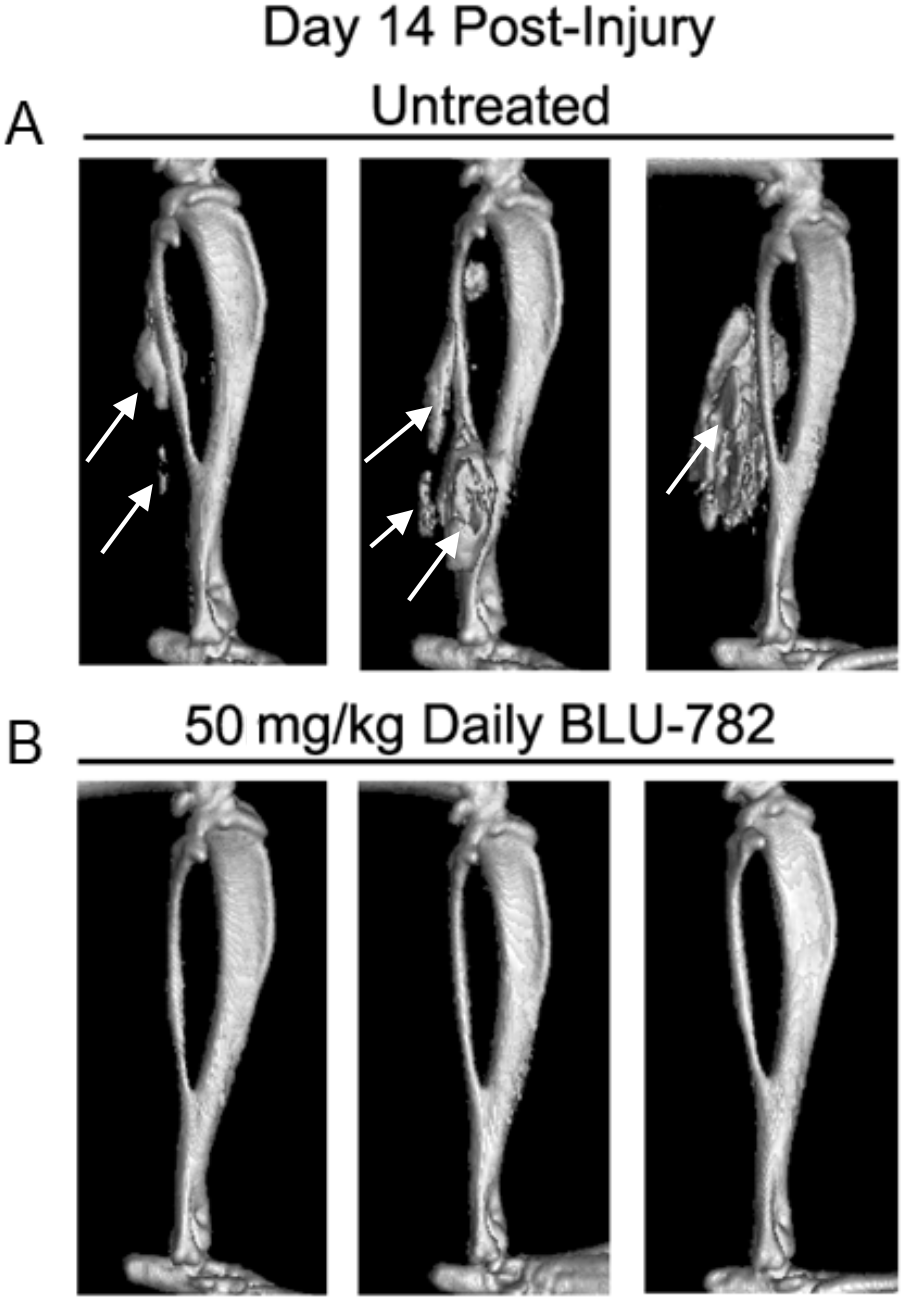
BLU-782 is highly effective at inhibiting injury-induced HO driven by R206H-FAPs in FOP mice. Representative µCT images of the lower hindlimbs of FOP mice (*Acvr1^tnR206H/+^*;*R26^NG/+^*;Tie2-Cre) taken 14 days after pinch injury of the posterior hindlimb muscle compartment. (A) Untreated FOP mice showing the range of HO volumes typically observed with this injury modality (n = 5). HO is shown at the arrows. (B) FOP mice were dosed daily with 50 mg/kg BLU-782 by oral gavage, beginning three days prior to injury through 13 dpi. HO was not observed (n = 6).

### Injury-induced global dysregulation of the R206H-FAP mRNA transcriptome

We performed RNA-seq of R206H-FAPs prior to muscle injury and at preskeletal lesional stages to identify genes that are dysregulated as R206H-FAPs are co-opted for pathological endochondral bone formation. Recombination of the conditional *Acvr1^tnR206H^* allele(^24–27^) was directed by Tie2-Cre, as injury-induced HO is highly penetrant with this Cre driver(^26^), and young adults do not develop spontaneous HO(^24^), which could have complicated the analysis. The *Acvr1^tnR206H^* allele allows discrimination of unrecombined and recombined cells based on the presence and absence of tdTomato fluorescence, respectively(^24^). This allele design feature allows collection of only recombined cells, which was particularly important for the present analysis since the Tie2-Cre-dependent recombination efficiency of the *Acvr1^tnR206H^* allele in FAPs of uninjured muscle is only approximately 10%(^24^).

2-3-month-old female FOP mice carrying the Cre-dependent GFP allele, *R26^NG^*(^38^) (*Acvr1^tnR206H/+^*;*R26^NG^*^/+^;Tie2-Cre), were pinch injured as above and all muscle tissue of the posterior lower hindlimb compartment was collected at 0, 1, 3, and 5 dpi. (Fig. S2). These time points encompass the early stages of lesional growth prior to the appearance of histologically identifiable cartilage, which is evident by 6-7 dpi(^24^). Parallel groups of mice received either a single subcutaneous injection of 10 mg/kg ActA-mAb on the day of injury, or daily oral dosing with 50 mg/kg BLU-782 beginning 3 days prior to injury (Fig. S2). Results were compared to untreated littermate control mice (WT for both *Acvr1* alleles), and to control mice treated with drug and collected at 5 dpi to determine how these treatments impacted the FAP transcriptome under normal regenerative conditions. FAPs from control mice, and recombined, R206H-FAPs from FOP mice were isolated by fluorescence-activated cell sorting as CD31-CD45-PDGFRα+SCA-1+GFP+tdTomato-cells, as previously described(^24–27^), and processed for RNA-seq using standard methods. Data is derived from three biological replicates for each condition.

Principle Component Analysis (PCA) was used to broadly assess the relatedness of mRNA transcriptomes between conditions. Notably, differential gene expression analysis using DESeq2 revealed that no genes were dysregulated in R206H-FAPs derived from untreated FOP mice at 0 and 1 dpi using the criteria of an adjusted p-value of < 0.05 and either a 50% increase or a 33% reduction in normalized counts. This indicates that FAPs are not transcriptionally primed for endochondral bone formation by expression of ACVR1(R206H) during development or juvenile stages. At 3 dpi, however, PCA showed that untreated R206H-FAPs diverged from both WT FAPs and R206H-FAPs treated with either drug (Fig. 2A). At 3 and 5 dpi, many genes were dysregulated in untreated R206H-FAPs, the majority of which were downregulated (Fig. 2B, E, H). We note that R206H-FAPs derived from one untreated FOP mouse at 5 dpi exhibited only modest gene expression changes. As the quantity of HO at endpoint can vary widely, this sample may have been derived from a limb destined for a comparatively minor HO response. Thus, the calculated number of significant gene expression changes at 5 dpi may be an underrepresentation, as all samples were included in statistical analyses.

**Figure 2.**
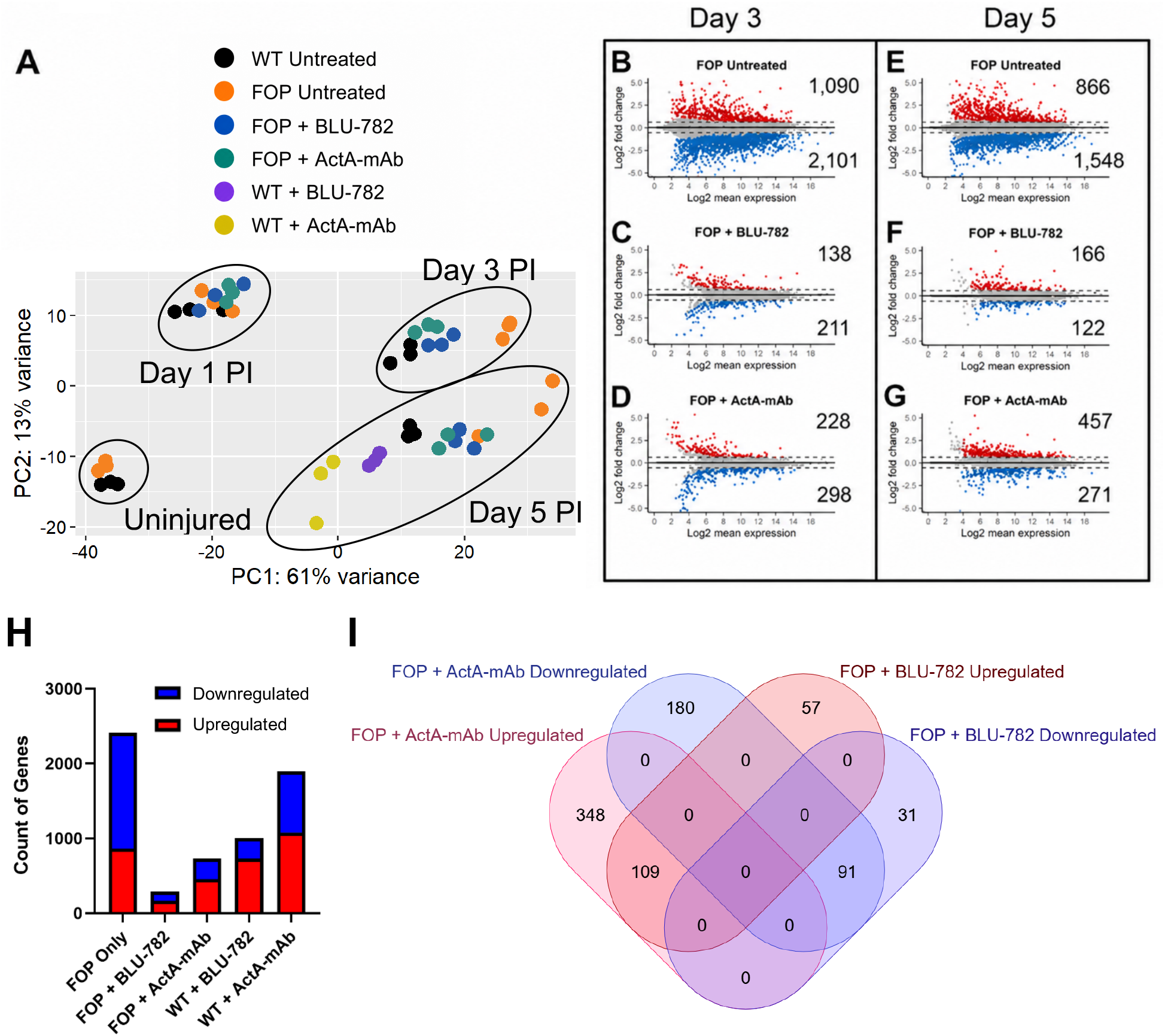
BLU-782 and ActA-mAb rescue the majority of dysregulated genes in R206H-FAP. (A) Principal Component Analysis (PCA) of all samples to assess relatedness within and among comparison groups. Each condition was repeated in triplicate. (B-G) MA plots showing log2 fold changes of genes expressed at 3 and 5 dpi in R206H-FAPs compared to FAPs derived from injured muscle of control mice (WT for *Acvr1*), plotted against log2 mean expression levels. Untreated and treated FOP mice are compared. Dotted lines represent ±0.585 log2 fold thresholds (50% upregulated or 33% downregulated) to add stringency to an adjusted p-value < 0.05 by the Wald test with Benjamini-Hochberg correction. Red and blue dots represent up- and downregulated genes, respectively. Gray dots represent genes that were not significant changed. Numbers of up- and downregulated genes are given. (H) Count of upregulated (red) and downregulated (blue) genes relative to WT controls at 5 dpi. (I) Venn diagram comparing dysregulated genes in R206H-FAPs derived from FOP mice treated with either BLU-782 or ActA-mAb and collected at 5 dpi.

Both ActA-mAb and BLU-782 substantially reduced the number of genes dysregulated in R206H-FAPs at both 3 and 5 dpi (Fig. 2B-I). Using FAPs derived from control regenerating muscle as the basis for comparison, treatment of FOP mice with BLU-782 rescued 89.1% and 88.1% of genes dysregulated in mutant FAPs at 3 and 5 dpi, respectively (Fig. 2B, C, E, F, H). Treatment of FOP mice with ActA-mAb at the time of injury corrected 83.5% (3 dpi) and 69.8% (5 dpi) of dysregulated genes (Fig. 2B, D, E, G, H). Comparative analysis of genes not corrected by drug treatment (Fig. 2I) and their relationship to dysregulated genes in drug-treated WT mice is presented in sections below.

### R206H-FAPs initiate skeletogenic gene expression by 3 days post injury

To assess the timing of gene expression associated with skeletogenic fates, we examined expression of chondrogenic and osteogenic genes over the post-injury time course. Interestingly, chondrogenic gene expression was already elevated at 3 dpi (Fig. 3G), 3-4 days before the appearance of histologically identifiable cartilage in this model. Chondrogenic genes include *Col2a1, Acan, Fgfr3, Ptch1, Chst8,* and *Chst11*, all of which are known transcriptional targets of SOX9, a master regulator of chondrogenesis(^39^). However, *Sox9* expression was not restricted to pre-cartilage stages or to R206H-FAPs; both FAPs from WT muscle and R206H-FAPs expressed *Sox9* prior to injury and at all post-injury time points (Fig. 3A, D; data not shown). While expression in WT FAPs may reflect their competency for skeletogenic differentiation, which has been demonstrated in some experimental settings(^12,17,40^), we note that *Sox9* expression is not restricted to the skeletogenic lineage, and it serves many functions at both embryonic and adult stages(^41^). Perhaps surprisingly, *Sox9* mRNA abundance was comparable in R206H-FAPs and WT FAPs at 3 dpi (Fig. 3A). However, as *Sox9* expression levels in R206H-FAPs destined for cartilage formation likely represent only a minority of total R206H-FAPs isolated from the entire posterior hindlimb compartment, measured levels of *Sox9* and other skeletogenic markers are likely an underrepresentation of mRNA levels of R206H-FAPs actively engaged in skeletogenesis. Despite this dilution effect, *Sox9* mRNA abundance was significantly elevated in R206H-FAPs by 5 dpi (Fig. 3D).

**Figure 3.**
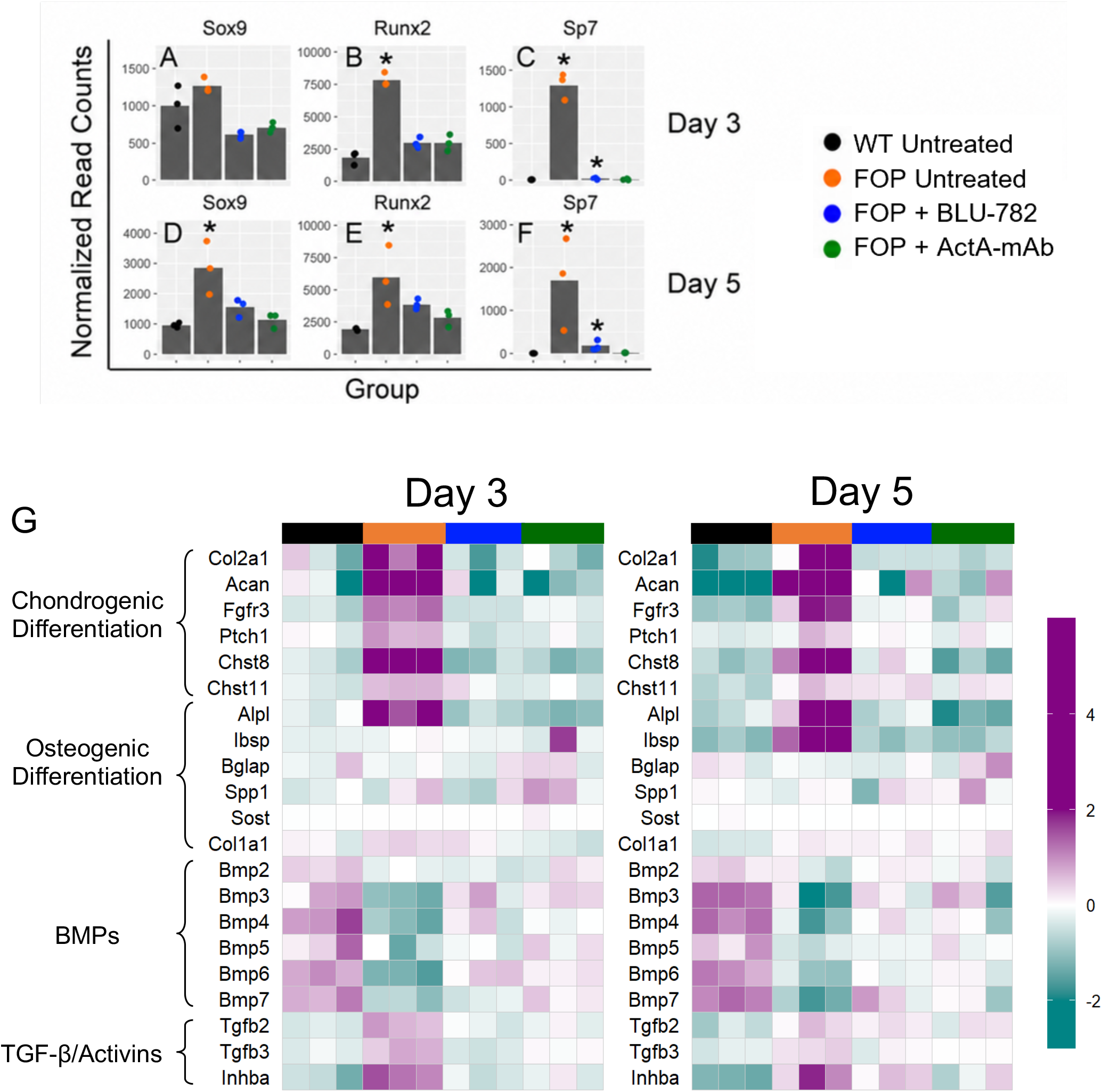
R206H-FAPs express chondrogenic and osteogenic genes at 3 and 5 dpi. (A-F) Bar graphs represent mean normalized counts of mRNAs encoding key transcriptional regulators of chondrogenesis and osteogenesis, *Sox9* (A, D), *Runx2* (B, E), and *Sp7* (osterix; C, F) at 3 and 5 dpi. (G) Heatmap of regularized log counts of markers of chondrogenic and osteogenic differentiation, BMPs, and TGF-β/activin ligands, at 3 and 5 dpi. Asterisks indicate significance compared to WT, as determined by an adjusted p-value < 0.05 and either 50% higher or 33% lower mean expression. The heatmap color scale indicates log2 values representing the difference with respect to the mean log2-transformed counts.

Bone is not radiographically evident until approximately 10 days after injury in this FOP mouse model (unpublished observations). Interestingly, mRNAs for the master regulators of osteoblast specification and differentiation, *Runx2* and *Sp7* (osterix; *Osx*), were already upregulated by 3 dpi (Fig. 3B, C). We note that *Runx2* and *Osx* also control late-stage chondrocyte differentiation(^42–44^), and their expression could reflect a chondrogenesis-specific function, assuming that these mRNAs are translated at this stage. Additionally, however, genes essential for bone mineralization are upregulated by 3 dpi (*Alpl*) and 5 dpi (*Alpl* and *Ibsp*) (Fig. 3G), indicating that the osteogenic transcriptional program had been initiated by at least some R206H-FAPs by these post-injury time points. Single-cell RNA-seq analyses will be required to define cell population structure at these times and determine whether R206H-FAP-derived cells that express cartilage or bone markers are already segregated into distinct subpopulations. Importantly, treatment with BLU-782 or ActA-mAb markedly downregulated both chondrogenic and osteogenic genes (Fig. 3A-G).

Genes encoding BMP and TGF-β ligands were also dysregulated. Many BMP-encoding genes were downregulated at both 3 and 5 dpi in R206H-FAPs, including *Bmp2*, *3*, *4*, *5*, *6*, and *7* (Fig. 3G). This group includes most of the osteogenic BMPs, suggesting that early FOP pathogenesis is not potentiated through increased activity of osteogenic BMPs, at least through FAP-derived autocrine or paracrine signaling mechanisms. Indeed, the strict dependency on activin A(^24,29^), and the observation that over-expression of ACVR1 does not sensitize FAPs to cognate BMP ligands(^26^), also argue against a primary role for osteogenic BMPs in early FOP pathogenesis. We also note that FAPs are highly adipogenic(^4,5,8^) and that BMP2 and BMP4 can induce commitment of pluripotent progenitors to an adipogenic fate(^45^). Thus, downregulation of these BMPs may serve to suppress the adipogenic fate early in the progression of FOP disease progression. *Tgfb2* and *Tgfb3* were both upregulated in R206H-FAPs at 3 and 5 dpi (Fig. 3G). Both ligands promote chondrogenesis in limb bud mesenchyme micromass cultures(^46^), and a TGF-β neutralizing antibody attenuated HO in an FOP mouse model(^47^). Interestingly, *Inhba*, the gene that encodes the monomer of activin A, was significantly upregulated in R206H-FAPs at 3 and 5 dpi (Fig. 3G; Fig. S3A, C). *Acvr1* was also significantly upregulated at 5 dpi (Fig. S3D). These results are consistent with a shift toward greater TGF-β/activin signaling in R206H FAPs in response to muscle injury. Upregulation of these ligands may create a positive feedback loop in which lesional growth is sustained and potentiated in an R206H-FAP-autonomous manner.

### Canonical Wnt and retinoic acid signaling programs are downregulated in R206H-FAPs

Canonical Wnt signaling has cell- and stage-specific effects on osteogenesis and adipogenesis, and FAPs are the main producers of Wnt ligands among skeletal muscle-resident cell populations(^48^). Generally, Wnt signaling promotes osteogenic differentiation and suppresses adipocyte differentiation, whereas at earlier developmental stages, Wnt signaling promotes stem cell maintenance and expansion(^49^). Interestingly, overrepresentation analysis (PANTHER) identified the gene ontology (GO) term, “Negative Regulation of Canonical Wnt Signaling” (GO:0090090). The genes encoding the canonical Wnt ligands WNT10B and WNT2 were downregulated at both 3 and 5 dpi. Importantly, in addition to Wnt pathway constituents, β-catenin target genes were downregulated at both 3 and 5 dpi; these genes include *Mmp2*, *Satb1*, and *Enpp2* (Fig. 4A). Upregulated genes include *Sfrp1* and *Sfrp2*, which encode secreted Frizzled-related inhibitors that act as soluble Wnt ligand traps. Additionally, they have been shown to bind β-catenin and inhibit its transcriptional activity(^50^). Finally, the non-canonical Wnt, WNT5A, which can antagonize canonical Wnt signaling by promoting the degradation of β-catenin(^51^), was upregulated at 3 dpi (Fig. 4B). The inhibitory effects of WNT5A are mediated by *Siah2*(^51^), which was upregulated at 5 dpi, following the period of transient *Wnt5a* upregulation (Fig. 4B, E). Collectively, these results suggest that altered Wnt signaling in R206H-FAPs is associated with their transcriptional reprogramming toward skeletogenic fates in FOP early lesional tissue.

**Figure 4.**
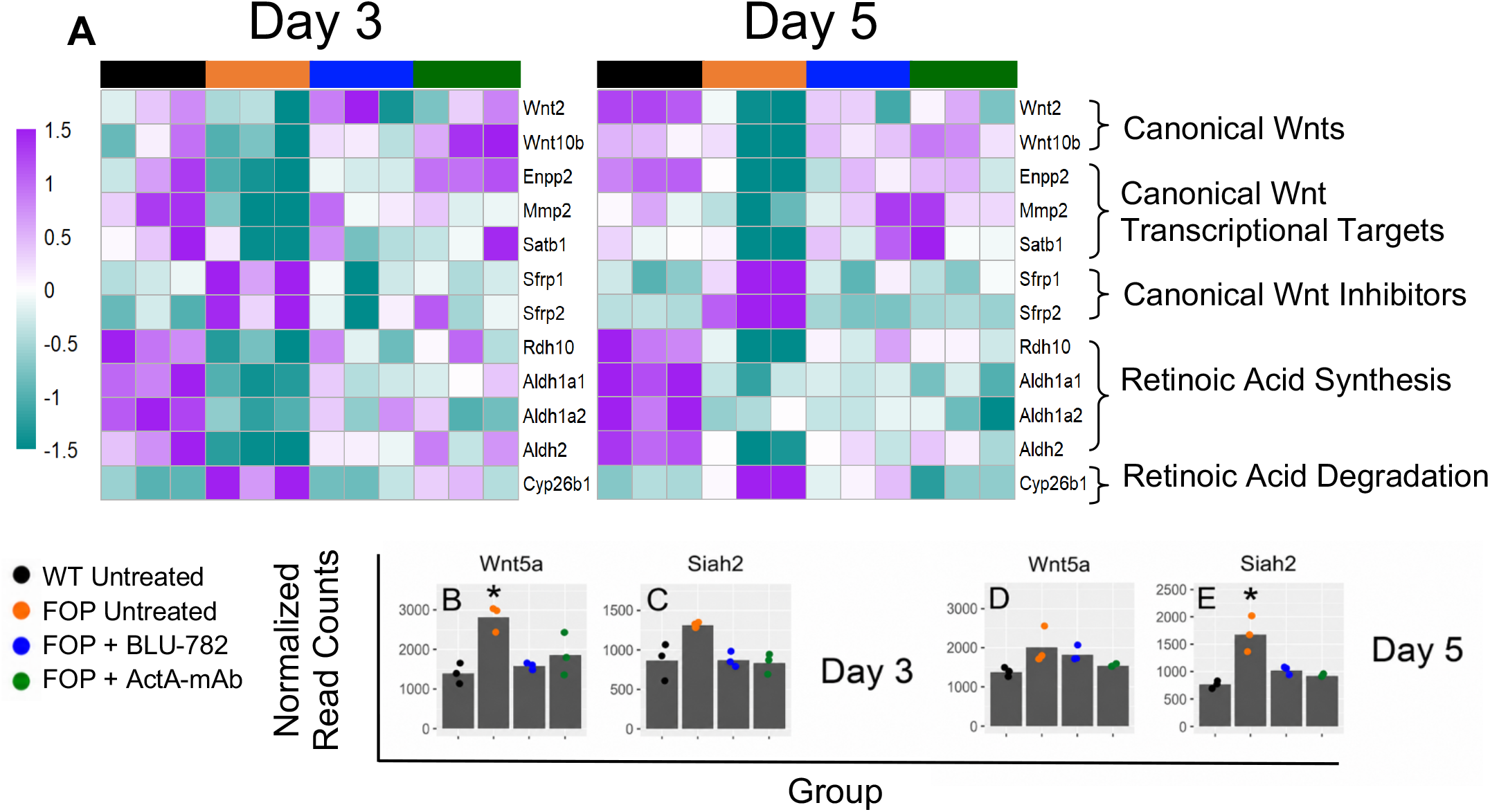
Downregulation of canonical Wnt and retinoic acid signaling in R206H-FAPs. (A) Heatmap of regularized log counts of genes exhibiting significant dysregulation in R206H-FAPs at 3 and 5 dpi. (B-E) Bar graphs representing the mean normalized counts of the non-canonical Wnt, *Wnt5a* (B, D), and its downstream target, *Siah2* (C, E), at 3 and 5 dpi. Asterisks indicate significance compared to WT, as determined by an adjusted p-value < 0.05 and either 50% higher or 33% lower mean expression. The heatmap color scale indicates log2 values representing the difference with respect to the mean log2-transformed counts.

Retinoic acid (RA) signaling inhibits chondrogenesis(^52,53^) and retinoic receptor agonists reduce HO in BMP induction models(^54^) and in mouse models of FOP(^27,55,56^). Notably, genes encoding the enzymes that catalyze the two-step conversion of retinol to RA (*Rdh10, Aldh1a1*, *Aldh1a2*, and *Aldh2*)(^57^) were downregulated at 3 and 5 dpi (Fig. 4A). Further, *Cyp26b1*, which encodes an enzyme that converts RA to inactive RA metabolites(^57^), was upregulated approximately four-fold at both 3 and 5 dpi (Fig. 4A). Collectively, these gene expression changes indicate a shift towards reduced RA signaling activity. As retinoic acid signaling has also been demonstrated to up-regulate canonical Wnt ligands and cause increased transcription of Wnt targets in FAPs(^10^), decreased RA signaling in R206H-FAPs may contribute to a decrease in canonical Wnt signaling. RA response genes that may influence R206H-FAP population growth dynamics were also dysregulated (see below). Treatment with BLU-782 or ActA-mAb was partially effective at normalizing expression of canonical Wnt and RA pathway genes (Fig. 4).

### Genes that control population growth are dysregulated in R206H-FAPs

We previously showed that R206H-FAPs exhibit activin A-dependent enhanced engraftment and population growth following transplantation(^24,27^). We used flow cytometry to determine whether R206H-FAP population growth after injury is enhanced in FOP mice relative to WT FAPs of control mice across the same time course used for RNA-seq analysis. While the number of R206H-FAPs and WT FAPs were similar at 3 dpi, R206H-FAPs were over two-fold more abundant by 5 dpi (Fig. 5A). This increase was likely an underrepresentation in areas of lesion formation, given the dilution effect resulting from the collection of FAPs from the entire posterior compartment of the lower hindlimb. Notably, treatment with either BLU-782 or ActA-mAb restored normal FAP population growth of R206H-FAPs (Fig. 5A).

**Figure 5.**
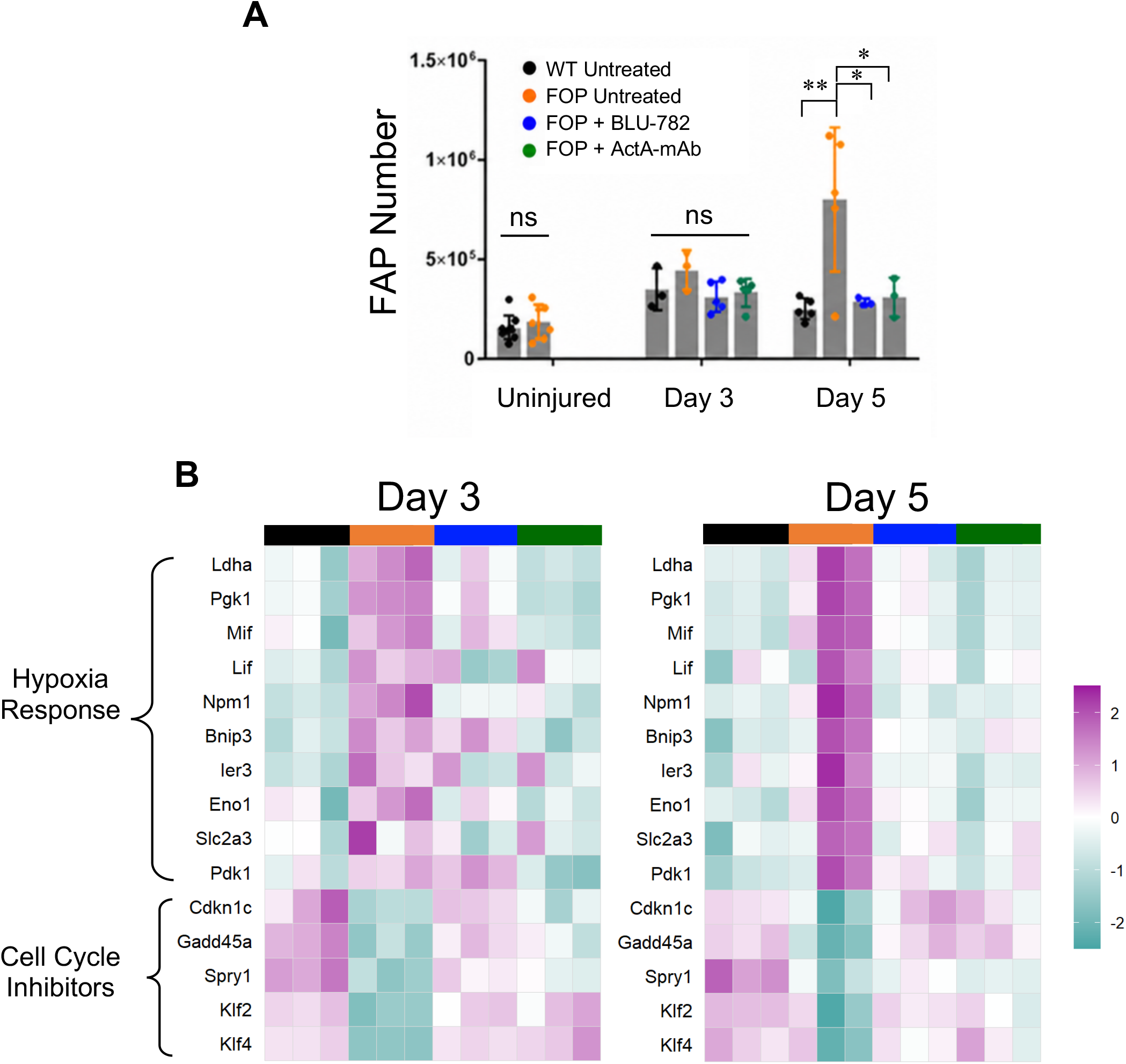
Dysregulation of hypoxia response genes, cell cycle regulators, and R206H-FAP growth in injured muscle of FOP mice. (A) Flow cytometry with counting beads was used to calculate FAP numbers harvested at the indicated post-injury time points. Error bars represent ± SD. Significance was determined using a one-way ANOVA with Tukey’s multiple comparison test. * p-value < 0.05, ** p-value < 0.01. (B) Heatmap of the regularized log counts of upregulated hypoxia response genes and downregulated inhibitors of the cell cycle at 3 and 5 dpi. Up- and downregulation was determined by an adjusted p-value < 0.05 and either 50% higher or 33% lower mean expression. The heatmap color scale indicates log2 values representing the difference with respect to the mean log2-transformed counts.

Two additional transcriptional targets of RA signaling that were downregulated in R206H-FAPs at both 3 and 5 dpi are tissue transglutaminase (*Tgm2*) and hypermethylated in cancer-1 (*Hic1*) (Fig. S4B, C, E, and F). *Tgm2* is a positive mediator of apoptosis in different tissues, including the interdigital mesenchyme of the developing limb(^58^), and it will be important to determine whether its downregulation plays a role in the decrease in FAP apoptosis observed in FOP mice 5 days after muscle injury(^59^). *Hic1* is a known regulator of quiescence in FAPs, as conditionally knocking out *Hic1* caused an increase in proliferative FAPs in the absence of an injury stimulus(^60^). In addition to these RA-responsive genes, overrepresentation analysis identified several differentially expressed genes involved in regulation of cell proliferation at 3 dpi, before R206H-FAP numbers were abnormally elevated at 5 dpi (Fig. 5A, B). Negative regulators of the cell cycle that were downregulated in R206H-FAPs at 3 dpi include *Gadd45a*, *Cdkn1c*, *Spry1*, *Klf2*, and *Klf4* (Fig. 5B), and these genes remained significantly downregulated at 5 dpi (Fig. 5B). By contrast, immediate early response 3 gene (Ier3), which positively regulates human adipose progenitor cell proliferation and renewal(^61^), was upregulated by 3 dpi (Fig. 5B). Collectively, altered regulation of mediators of apoptosis and the cell cycle could contribute to enhanced R206H-FAP population growth, which precedes the appearance of definitive cartilage and is correlated with HO severity(^27^).

### Hypoxia response genes are upregulated in R206H-FAPs

Formation of precartilage condensations require a hypoxic cellular environment, and in FOP mouse models, hypoxia promotes HO while the inhibition of hypoxia prevents HO(^62,63^). Several hypoxia response genes were upregulated in R206H-FAPs (Fig. 5B). *Hif1a*, which encodes a master regulator of the hypoxia response, was upregulated in R206H-FAPs derived from injured muscle (data not shown). However, *Hif1a* was not classified as dysregulated because *Hif1a* was also upregulated in FAPs isolated from injured muscle of control mice. Hypoxia response genes categorized as dysregulated at 3 and 5 dpi include *Ier3*, the glycolytic enzyme-encoding genes *Pgk1* and *Ldha*, *Bnip3*, which encodes a mitophagy inducer(^64^), and the genes for the cytokines LIF and MIF. At 5 dpi, two additional genes that encode glycolytic enzymes (PDK1 and ENO1) were upregulated. *Mif* is a transcriptional target of HIF1-α(^65^), and in turn, MIF can modulate HIF1-α activity(^66^). HIF1-α is required for chondrocyte survival(^67^) and differentiation(^68^), and both MIF and HIF1-α are direct regulators of *Sox9*(^68,69^). *Inhba*, which was upregulated at 3 and 5 dpi (Fig. 3G; Fig. S2A, C), is also a hypoxia response gene in some contexts; *INHBA* was induced in human articular chondrocytes under hypoxic conditions and induction was dependent on HIFs but not SOX9(^70^). Interestingly, expression of *Ier3* was enhanced by activin A(^61^), suggesting that activin A may both respond and contribute to the hypoxia-induced transcriptional program. Inhibiting ACVR1 signaling with BLU-782 or blocking the function of activin A with ActA-mAb prevented the upregulation of hypoxia responsive genes in FOP mice (Fig. 5B).

### R206H-FAPs alter their environment through dysregulated production of ECM components and ligands that mediate cell-cell interactions

ECM constituents were broadly downregulated in R206H-FAPs. Of 59 dysregulated ECM structural constituent genes (GO:0005201), 48 were downregulated (Fig. 6A). These downregulated ECM genes include various laminins, collagens, and nidogens used to construct the basement membrane of skeletal muscle and vasculature(^71^). Also broadly downregulated were genes encoding integrins (GO:0033627) (Fig. 6A), broadly expressed receptors that regulate myriad processes, including cell migration, proliferation, differentiation, and cell survival(^72,73^). While the functional consequences of diminished integrin-ECM interactions require further investigation, these gene expression changes indicate that R206H-FAPs alter their environment, which in turn alters their behavior and that of neighboring cell types. In this context, FAPs have been shown to serve positive effector functions in regeneration by interacting with satellite cells and influencing myogenic differentiation(^14,74,75^). We also examined data derived from the scRNA-seq study of injured muscle conducted by De Micheli et al.(^76^), which included ligand-receptor interactions at 5 and 7 days after notexin-induced muscle injury (see their Figure S6). Among the 33 highest scoring ligand-myogenic cell receptor interactions, 14 were downregulated and 5 were upregulated in R206H-FAPs compared to FAPs from regenerating WT muscle (Fig. 6B). Among the downregulated genes were the ECM constituents *Myoc*, *Dcn*, and laminin subunit, *Lamc2*. *Myoc* and *Dcn* can promote differentiation and hypertrophy of newly differentiated myotubes in C2C12 myogenic cell culture(^77^). As regeneration is impaired when *Acvr1^R206H^* expression is targeted to FAPs(^24,59,78^), these data suggest that interactions between FAPs and satellite cells that normally promote regeneration are disrupted in FOP mice.

**Figure 6.**
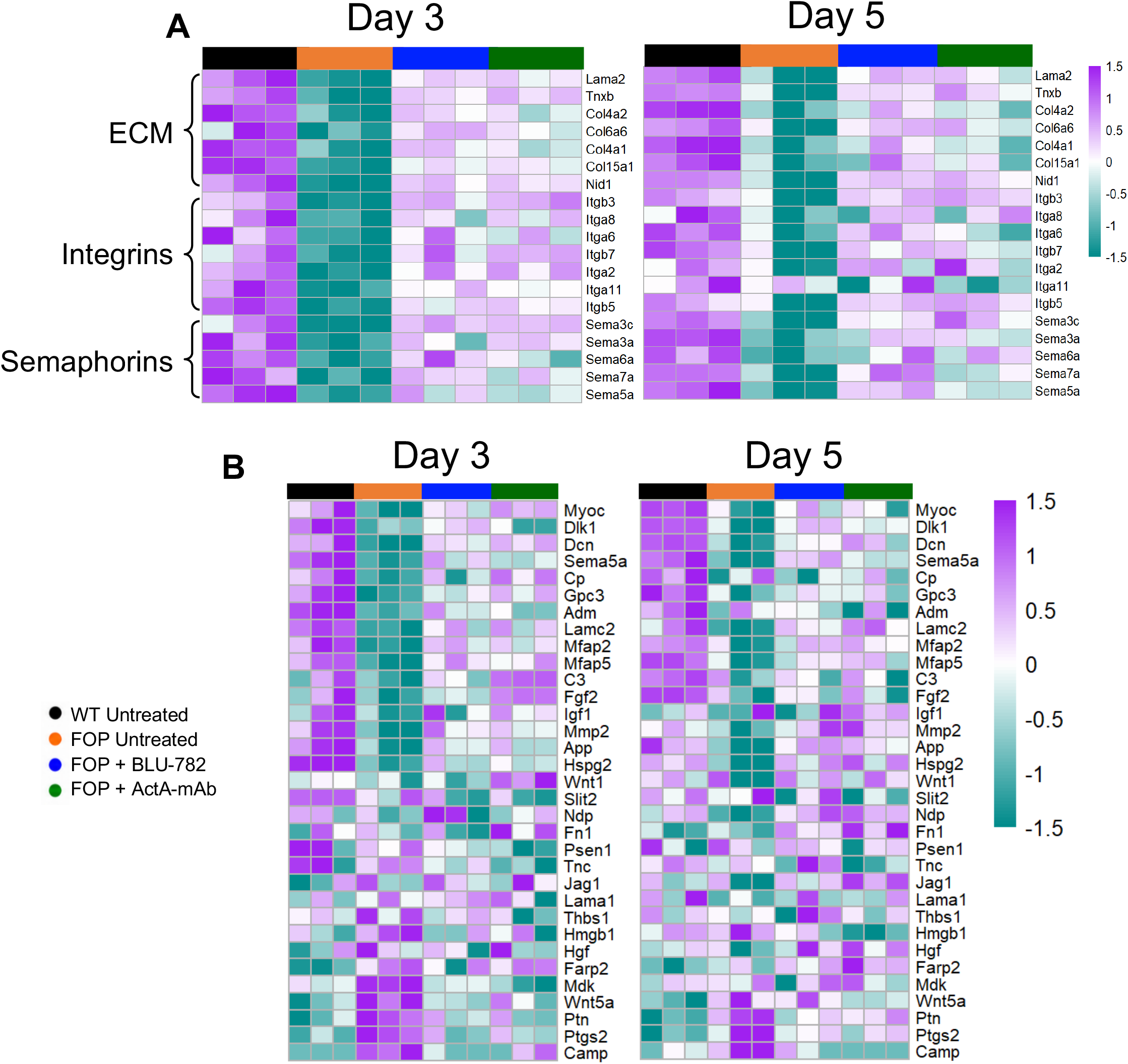
ECM constituents and mediators of signaling with other cell types are downregulated in R206H-FAPs at 3 and 5 dpi. (A) Heatmap of regularized log counts at 3 and 5 dpi of downregulated ECM structural constituents, integrins, and semaphorins. (B) Heatmap of 33 FAP ligands with the highest scores on ligand-muscle stem cell receptor interactions at 5 and 7 days after muscle injury, as determined by De Micheli et al.(^77^). The heatmap color scale indicates log2 values representing the difference relative to the mean of the log2-transformed counts.

### Inhibition of activin A and ACVR1 activity have overlapping but distinct effects on the transcriptome of mutant FAPs

BLU-782 and ActA-mAb treatment corrected overlapping sets of genes; however, many genes remained dysregulated (Fig. 2B-G, H, I). This analysis focused on 5 dpi, as this time point showed substantial divergence between treatments in apparent efficacy in restoring the R206H-FAP transcriptome (Fig. 2B-G, H, I). Genes that were dysregulated at 5 dpi with drug treatment comprise three classes based on whether both ActA-mAb and BLU-782 or only one drug failed to normalize gene expression (Fig. 2H, I; Fig. 7A-D). GO analysis of genes that were uncorrected by both ActA-mAb and BLU-782 at 5 dpi identified biological processes that were largely involved in cell signaling, development, and differentiation (Fig. 7A, B). Transcription factors and signaling molecules that remained dysregulated in both treatment groups include *Bmp4*, *Bmp6*, *Pi16*, *Nfatc2*, and *Dlx5*, among several others. A larger number of genes remained dysregulated in ActA-mAb-treated FOP mice compared to treatment with BLU-782 (Fig. 2H, I, Fig. 7A). Genes that remained uncorrected in R206H-FAPs only when treated with ActA-mAb comprise GO terms involved in biological processes that include hypoxia, signaling, and cell-cell communication (Fig. 7C). In contrast, GO analysis of genes that were uncorrected in R206H-FAPs only when treated with BLU-782 revealed terms involved in blood circulation and control of vessel diameter (Fig. 7D).

**Figure 7.**
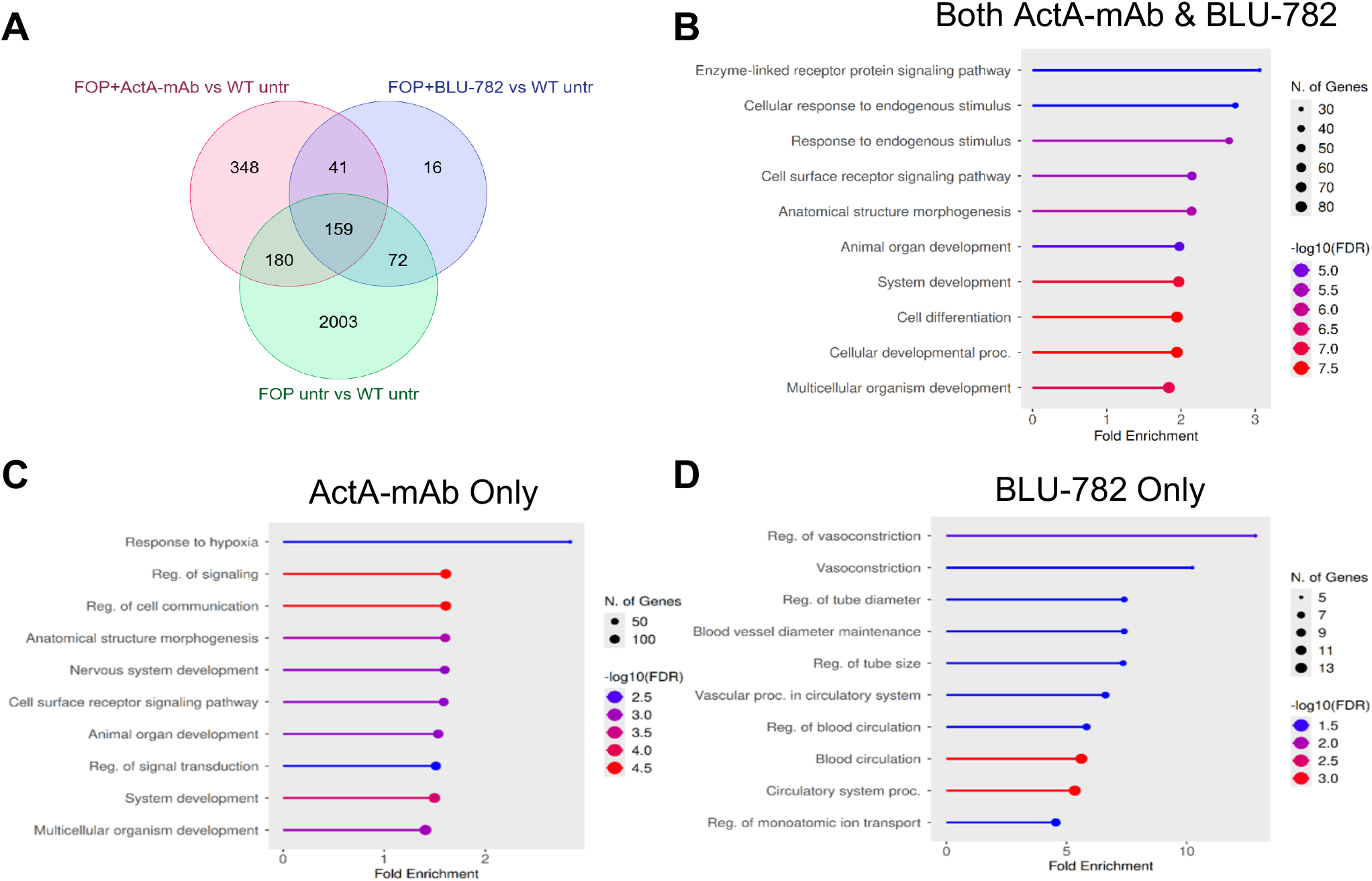
Comparison of dysregulated genes and processes in R206H-FAPs derived from FOP mice treated with either ActA-mAb or BLU-782. Analyses were conducted using data from 5 dpi. (A) Venn diagram showing the relationship between dysregulated genes in the two drug treatment groups and in untreated FOP mice. (B-D) Lollipop plots of the genes impacted by ActA-mAb and BLU-782 treatment. The top 10 Gene Ontology Biological Processes with FDR < 0.05 and a minimum of 10 genes per pathway are shown. Processes affected by both treatments (B), ActA-mAb only (C), or BLU-782 only (D), are shown.

### Dysregulation of gene expression by ActA-mAb and BLU-782 in FAPs derived from injured muscle of control mice

We next evaluated how each drug affected the FAP transcriptome of control mice (WT for *Acvr1*) 5 days after muscle injury. The number of dysregulated genes and presumptive pathways affected was greater in ActA-mAb-treated control mice (Fig. 2H; Fig. 8A-D). 1,898 genes were significantly dysregulated in ActA-mAb treated controls, compared to 2,414 dysregulated genes in R206H-FAPs derived from untreated FOP mice (Fig. 2H; Fig. 8A). A substantial but smaller number of genes were dysregulated in BLU-782-treated control mice (1,004 genes; Fig. 2H; Fig. 8A). Of the total number of dysregulated genes, 643 genes (28.5%) were affected by both treatments, 1,255 genes (55.6%) were affected only by ActA-mAb treatment, and 361 genes (16%) were dysregulated only by treatment with BLU-782 (Fig. 2H; Fig. 8A).

**Figure 8.**
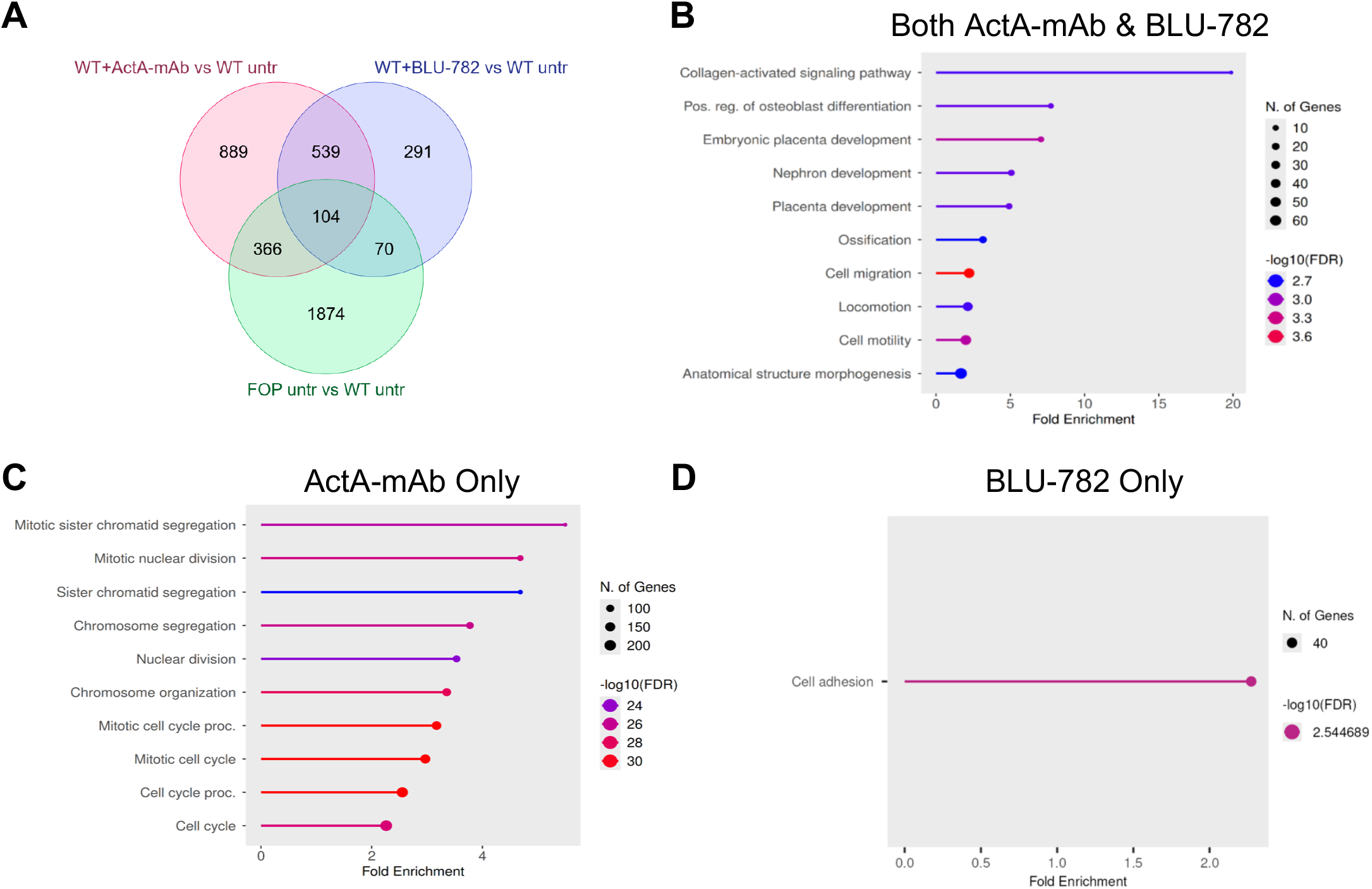
Comparison of dysregulated genes and processes in FAPs derived from WT mice treated with either ActA-mAb or BLU-782. Analyses were conducted using data from 5 dpi. (A) Venn diagram of dysregulated genes comparing ActA-mAb and BLU-782 treatment of WT mice, and their relationship to dysregulated genes in R206H-FAPs derived from untreated FOP mice. (B-D) Lollipop plots of the processes impacted by both drugs (B), ActA-mAb only (C), or BLU-782 only (D). The top 10 Gene Ontology Biological Processes with FDR < 0.05 and a minimum of 10 genes per pathway are shown.

Of the genes affected by both treatments in control mice, GO analysis identified biological processes terms involved in collagen-activated signaling, osteoblast differentiation, and ossification (Fig. 8B). The primary GO biological processes impacted by ActA-mAb treatment that were not dysregulated in BLU-782-treated control mice include a number of genes involved in the cell cycle, and mitotic division (Fig. 8C). GO analysis of genes that were only dysregulated in BLU-782-treated control mice revealed only one biological process, cell adhesion (Fig. 8D).

The incomplete normalization of gene expression in R206H-FAPs after drug treatment could reflect inefficiencies in drug action (i.e., incomplete inhibition of ACVR1(R206H)-dependent pathogenic signaling) (Fig. 9A, E). Alternatively, some genes could have remained uncorrected even with efficient inhibition of ACVR1(R206H) activity if the drugs affected gene expression downstream of normal activin A or ACVR1 signaling (Fig. 9A, D). Further, expression of some genes could be corrected even with insufficient inhibition of pathogenic signaling in cases where the drug caused gene dysregulation in the opposite direction (i.e., up- vs downregulation) as that caused by ACVR1(R206H) activity (Fig. 9C). To assess whether drug effects might have contributed to the classification of genes as corrected or uncorrected, we directly compared dysregulated genes in FOP-untreated, FOP-treated, and WT-treated mice (Fig. 9F, G).

**Figure 9.**
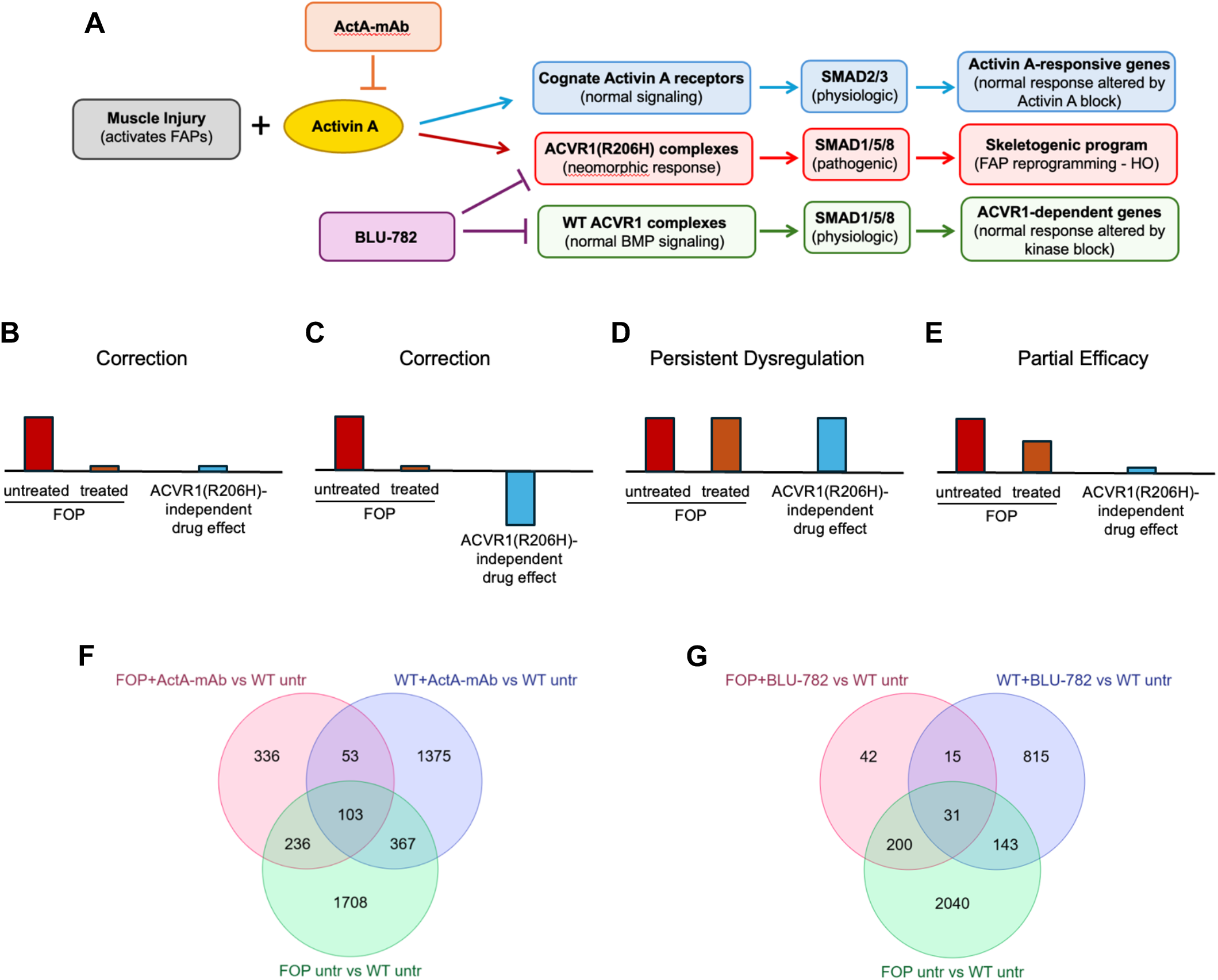
Comparison of dysregulated genes in FAPs derived from drug-treated WT and FOP mice and possible mechanisms that underpin drug effects. Analyses were conducted using data from 5 dpi. (A) Pathways affected by blocking activin A or inhibiting ACVR1/ACVR1(R206H) kinase activity. Note that assuming an equimolar amount of WT and mutant receptors in heterozygous cells (*Acvr1^R206H/+^*), ∼25% of receptor complexes should be WT for ACVR1 (see ref. 26). (B-E) Idealized examples of categories of genes based on how ACVR1(R206H)-independent effects of drug treatment and partial drug efficacy affect normalization of gene expression. The horizontal line in each panel represents normal expression of a given gene, as determined by RNA-seq of FAPs derived from injured WT muscle. (B) Expression of this group of dysregulated genes in R206H-FAPs is corrected because the drug is sufficiently effective in blocking ACVR1(R206H) pathogenic signaling and their expression is not affected by drug treatment in an ACVR1(R206H)-independent manner. (C) Some genes may be classified as corrected because of opposing effects on gene expression of ACVR1(R206H) pathogenic signaling and ACVR1(R206H)-independent signaling resulting from drug treatment. The figure does not distinguish directionality of changes in gene expression (upregulation vs downregulation). (D) Dysregulated gene expression may persist regardless of efficacy of drug action if the directionality of expression is the same as for ACVR1(R206H) pathogenic signaling. Note that a pronounced opposing effect of drug action could also result in persistent dysregulation (not depicted). (E) Some genes may remain dysregulated because the degree of inhibition of pathogenic signaling in the presence of drug is not sufficient to normalize expression of this gene class. R206H-independent effects may also contribute to expression levels (not depicted). (F, G) Venn diagrams showing dysregulated genes in ActA-mAb-treated (F) and BLU-782-treated (G) mice and their relationship to dysregulated genes in untreated FOP mice.

Among the 2,414 genes that were dysregulated in R206H-FAPs derived from untreated FOP mice, 103 genes were both uncorrected by treatment of FOP mice with ActA-mAb and were dysregulated in WT mice treated with drug (Fig. 9F; ∼14% of dysregulated genes in treated FOP mice). Some genes in this category would include those that were dysregulated in the same direction by ACVR1(R206H) activity and inhibition of activin A (Fig. 9D). Other genes in this category could include genes dysregulated in the opposite direction by drug treatment but to a greater and significant extent compared to that caused by ACVR1(R206H) activity (not shown). 31 uncorrected genes (∼11% of dysregulated genes in treated FOP mice) fell into this category in BLU-782-treated FOP mice (Fig. 9G). A set of 367 genes were corrected by ActA-mAb treatment of FOP mice but were dysregulated in ActA-mAb-treated WT mice (Fig. 9F). This class of genes (∼22% of corrected genes) is likely the consequence of opposing effects on gene expression of ACVR1(R206H) activity and activin A inhibition (Fig. 9C). In BLU-782-treated FOP mice, 143 genes comprised this category (∼7% of corrected genes; Fig. 9C, G). Presumably, ActA-mAb-dependent gene dysregulation largely reflects the blockade of activin A’s functional interactions with its cognate receptors, which normally activate signaling via SMAD2/3 (Fig. 9A). In contrast, BLU-782-dependent gene dysregulation is likely the consequence of blocking signaling through WT ACVR1-containing complexes (Fig. 9A). However, we cannot exclude the possibility that off-target effects contributed to drug-dependent gene dysregulation. Finally, we note that a proportion of uncorrected genes in treated FOP mice probably are the consequence of only partial drug efficacy in blocking pathogenic signaling through ACVR1(R206H)-containing complexes (Fig. 9A, E). This category includes genes dysregulated in both treated and untreated FOP mice but not dysregulated in WT mice (236 genes in ActA-mAb-treated mice, Fig. 9F; 200 genes in BLU-782-treated mice, Fig. 9G).

## DISCUSSION

FAPs are a transcriptionally heterogenous population of multipotent mesenchymal cells capable of differentiating into adipocytes, fibroblasts, cartilage, and bone(^4,5,12,24,79^). While a detailed understanding of the molecular and signaling pathways that regulate their adipogenic and fibrogenic capacity is emerging(^14^), little is known of the intrinsic factors and influences of the tissue environment that co-opt FAPs for pathological cartilage and bone formation. In the present study, we defined transcriptional changes in FAPs caused by expression of the FOP disease-causing gene, *Acvr1^R206H^*, to identify genes and pathways associated with injury-induced reprogramming of FAPs towards skeletogenic fates. The transcriptional effects of two leading therapeutic agents on FAPs derived from injured muscle of control and FOP mice were also defined.

Although *Acvr1^R206H^* is expressed in FAPs in the uninjured limb and at 1 dpi, effects of presumptive signaling via ACVR1(R206H) are not manifested at these stages; FAPs derived from both control and FOP mice showed similar changes in gene expression at this early stage after injury. Thus, although *Acvr1^R206H^* expression in cultured mouse embryonic fibroblasts poises them for osteochondral lineage progression(^80^), the present gene expression analysis is consistent with the view that R206H-FAPs are not transcriptionally pre-programmed for skeletal development prior to, or shortly after, muscle injury. We note, however, that developmental differences between WT and R206H-FAPs could be encoded epigenetically, a possibility that will require further investigation.

The current study identified candidate genes and putative pathways whose dysregulation in R206H-FAPs drives lesion formation, and inferences drawn here will be interrogated in future work. As lineage tracing studies have shown that Tie2+ FAPs do not build the normal skeleton(^15,24^), some of the dysregulated genes likely regulate critical pathways that may be uniquely responsible for early reprogramming of FAPs for pathological bone formation. By 3 dpi, R206H-FAPs displayed widespread dysregulation of gene expression, which included genes encoding signaling pathway components and transcription factors that regulate not only skeletal development, but also embryonic development and adult homeostasis. This identifies the stages between 1 and 3 dpi as a period of dramatic transcriptional reprogramming of FAPs. In BLU-782 efficacy studies, delaying dosing with this ACVR1 kinase inhibitor until 2 dpi blocked HO, while initiating dosing at 4 or 6 dpi still substantially reduced HO(^30^). Similarly, HO was inhibited when dosing with ActA-mAb was delayed until 3 dpi(^24^). Together, these data suggest that early transcriptional reprogramming of HO progenitors in FOP remains labile, requiring ongoing ACVR1(R206H) activity for their stable commitment to cartilage and bone fates.

Despite standardization of the anatomical location and method of pinch injury, the site of preskeletal lesional tissue formation could not be predicted with certainty, necessitating the isolation of R206H-FAPs from the entire posterior compartment of the lower hindlimb. Inclusion in the cell isolation procedure of tissue not actively engaged in lesion formation likely resulted in substantial “dilution” of the R206H-FAPs that were destined for skeletogenesis, dampening the magnitude of measured transcriptional changes. Because of these considerations, only female FOP mice were used because they display a significantly more robust injury-induced HO response than male mice in this model(^31^), thereby moderating the dilution effect. Reasons for this sex difference are not clear, although the female bias appears to be intrinsic to the cells themselves rather than being dictated by systemic factors(^31^). Parallel transcriptional studies in male FOP mice are needed to determine whether qualitative or quantitative differences in gene expression implicate particular genes, gene regulatory networks, or cellular processes. Obvious gene candidates include *Inhba* and *Acvr1/Acvr1^R206H^*, which are upregulated in R206H-FAPs. In a recent preclinical gene therapy study, Yang et al. showed that the *Acvr1^R206H^* and WT *Acvr1* alleles in heterozygous human FOP iPSCs were expressed at a ratio of approximately 2 to 1(^81^), and our work in FOP mice has shown that disease severity is highly influenced by the relative dosage of mutant and WT alleles(^24,26^). These data raise the possibility that sex differences in allele-specific receptor expression may contribute to qualitatively or quantitatively different transcriptional responses in male and female FOP mice. Even if these alleles are not expressed at different levels, an overall increase in expression of both ACVR1 and ACVR1(R206H) receptors in females could also produce a female bias, assuming that activin A ligand in the tissue environment is present in excess.

FAPs regulate muscle regeneration through multiple mechanisms that include immunomodulation at early post-injury times, conditioning of the ECM, and secretion of signaling molecules that affect satellite cell proliferation, differentiation, and fusion(^14^). Expression of *Acvr1^R206H^* in FAPs has the pathological effects of driving skeletogenesis in a cell-autonomous manner(^24^), disrupting muscle integrity(^78^), and impairing muscle regeneration cell-non-autonomously(^24,59,78^). Accordingly, the differentiation capacity of satellite cells was diminished when co-cultured with R206H-FAPs(^59^), an effect of ACVR1(R206H) activity that could result from dysregulated expression of positive and negative effectors of satellite cell functions. In this context, many genes that encode ECM constituents and ligands predicted to interact with satellite cells(^76,82,83^) were dysregulated, and future work is required to determine how these gene expression changes affect satellite cell processes. As FAPs express osteogenic genes following satellite cell ablation(^84^), impaired satellite cell function caused by ACVR1(R206H) activity in FAPs may, in turn, facilitate osteogenic lineage progression and differentiation of mutant FAPs in FOP mice.

An aspect of FOP pathogenesis that deserves increased attention is the effect of ACVR1(R206H) signaling on R206H-FAP population growth. Cartilage formation by competent progenitors is favored under conditions of high cell density and a hypoxic environment(^85,86^). The present analysis identified biological processes related to cell cycle regulation, apoptosis, hypoxia, and the ECM, dysregulation of which may impact R206H-FAP population growth cell-autonomously or cell-non-autonomously. Further, we identified several negative regulators of cell proliferation that were downregulated, including the RA-responsive gene *Hic1*(^60^). The RA transcriptional target *Tgm2*, which is a positive regulator of apoptosis in some contexts(^58^), was also downregulated. We previously showed using a cell transplantation model that population growth of R206H-FAPs is dependent on activin A(^27^). Here, R206H-FAP population growth was significantly greater than that of WT FAPs in control, regenerating muscle, as evidenced by the significantly elevated number of total R206H-FAPs at 5 dpi. Our study did not determine whether increased proliferation, decreased apoptosis, or both, were responsible for elevated R206H-FAP numbers. A previous study demonstrated that fewer FAPs undergo apoptosis in FOP mice after muscle injury(^59^), although whether this was a cell-autonomous effect resulting from *Acvr1^R206H^* expression in FAPs was not addressed. While the precise mechanisms require further investigation, it is noteworthy that both ActA-mAb and BLU-782 reduced R206H-FAP numbers to control levels, consistent with the corrective action of these drugs upstream of the major phase of skeletogenic differentiation.

Given the distinct mechanisms of action of ActA-mAb and BLU-782, it is not surprising that expression of distinct, but overlapping, sets of genes were normalized or remained uncorrected when FOP mice were treated with these therapeutic agents. Activin A typically signals by binding to its cognate receptor complexes and activating SMAD2/3. Combined with inhibition of ACVR1(R206H)-dependent signaling, inhibition of the activin A-driven SMAD2/3 signaling axis likely contributed to the transcriptional output of R206H-FAPs. Relatedly, BLU-782 is a highly specific inhibitor of ACVR1 kinase activity, but it does not selectively inhibit ACVR1(R206H), and R206H-FAP transcription in BLU-782-treated mice likely includes the effect of WT ACVR1 inhibition. Thus, the transcriptional output of R206H-FAPs derived from drug-treated FOP mice reflects both the efficacy of inhibiting ACVR1(R206H) pathogenic signaling and the consequence of ACVR1(R206H)-independent signaling activities of activin A and BLU-782. Gene expression analysis of FAPs from drug-treated control mice supports this conclusion and further suggests that the directionality of ACVR1(R206H)-independent gene expression changes can either mitigated or sustained the dysregulation of sets of genes in R206H-FAPs.

We note that despite gene dysregulation caused by the drugs themselves, and the substantial, but incomplete, normalization of ACVR1(R206H)-dependent gene expression, both treatments are highly effective at blocking HO in FOP mice(^24,29,30^) (present study). These observations argue that restoration to WT levels of expression of genes that remained dysregulated with treatment is not essential for blocking HO in FOP mice. However, we cannot rule out involvement of these uncorrected genes in processes associated with early lesion formation or other pathological processes upstream of definitive skeletal HO stages. In this regard, since muscle integrity is impaired in FOP mice(^78^), and muscle loss contributes to patient morbidity, it will be important to determine whether muscle integrity is maintained by these therapeutic agents.

During the course of this work, results of phase 2 and phase 3 clinical trials for FOP have been reported. The anti-activin A antibody garetosmab was highly effective at blocking new HO in the Phase 2 LUMINA-1 (NCT03188666)(^87^) and Phase 3 OPTIMA (NCT05394116) trials, and it recently received FDA approval for use in adults with FOP. Positive results were also reported for the highly selective ACVR1 inhibitor zilurgisertib (INCB000928) in the Phase 2 Progress trial (NCT05090891). In contrast, the Phase 2 FALKON trial (NCT05039515) evaluating fidrisertib (BLU-782) did not meet its primary endpoint of reducing new HO in individuals with FOP, and the study has been closed. Whether the disparate outcomes of ACVR1 kinase inhibition are due to differences in efficacy of inhibiting ACVR1(R206H) at the doses used or other aspects of drug action is unclear. Further interrogation of the genes that are normalized or remain uncorrected by each treatment may provide insights into the differential responses observed in humans.

## ACKNOWLEDMENTS

We thank members of the Goldhamer Lab for helpful input throughout the course of this work, Heather Jamieson for critical review of the manuscript, and Dr. Bo Reese (Scientific Director of the Center for Genome Innovation at the University of Connecticut) for RNA sequencing. The anti-activin A monoclonal antibody and BLU-782 were kindly provided by Acceleron Pharma and Blueprint Medicines Corporation, respectively. This work was funded by a grant from the NIH (R01AR072052) to DJG and by a sponsored research agreement between Blueprint Medicines and the University of Connecticut.

