## Supplemental Figures for "Pathogenic transcriptional reprogramming of fibro-adipogenic progenitors in mice with FOP and its mitigation by inhibition of ACVR1 and activin A"

Stoessel et al.

### SUPPLEMENTAL MATERIALS

#### Supplemental Figures 1-4

**Figure S1:** Examples of tibial exostosis after muscle injury in both BLU-782-treated and untreated FOP mice

**Figure S2:** Drug treatment schedule and bulk RNA-seq workflow

**Figure S3:** *Inhba* and *Acvr1* are upregulated in R206H-FAPs after muscle injury

**Figure S4:** R206H-FAPs upregulate a master adipogenic gene and downregulate retinoic acid targets after injury

**Supplemental Figure 1**

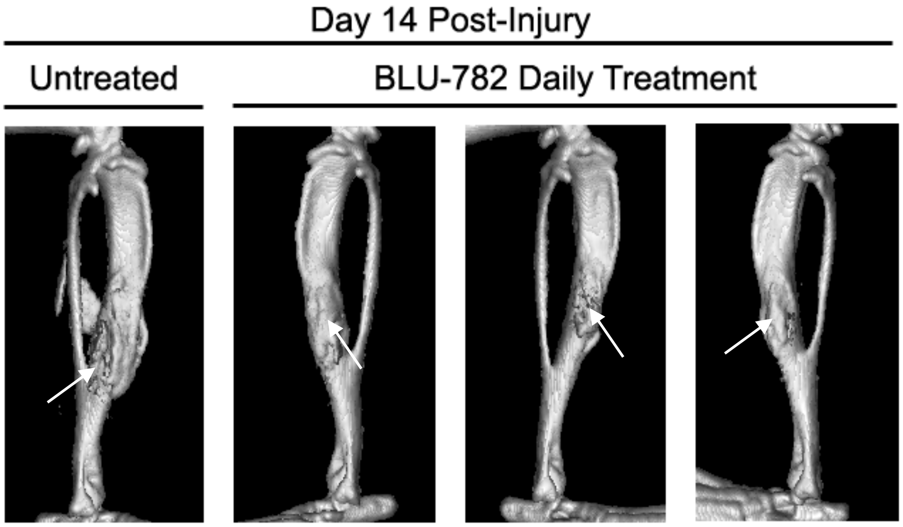


**Figure S1. Examples of tibial exostosis after muscle injury in both BLU-782-treated and untreated FOP mice.** Untreated and treated FOP mice (*Acvr1*$^{tnR206H/+}$;*R26*$^{NG/+}$;Tie2-Cre) were imaged by μCT at 14 dpi. Both hindlimbs of each mouse were injured. 3 of 12 hindlimbs (3 of 6 mice) of BLU-782-treated FOP mice developed small tibial overgrowths. 3 of 10 hindlimbs (3 of 5 mice) of untreated FOP mice also showed tibial overgrowths, which were sometimes obscured by closely associated intramuscular HO. Exostosis is shown at the arrows.

### Supplemental Figure 2

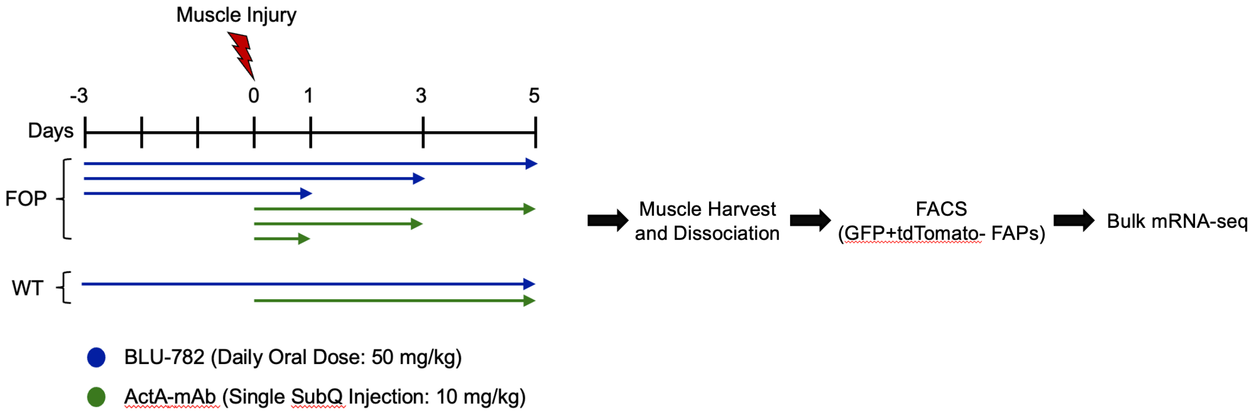


**Figure S2. Drug treatment schedule and bulk mRNA-seq workflow.** Control (*R26*$^{NG/+}$;Tie2-Cre) and FOP (*Acvr1*$^{tnR206H/+}$;*R26*$^{NG/+}$;Tie2-Cre) mice either received no treatment or were treated by the indicated dosing regimens with either BLU-782 or ActA-mAb. Muscles of the lower hindlimb posterior compartment were pinch-injured, and muscles collected at the indicated post-injury time points (n = 3; 2-3-month-old females). Control mice (WT for *Acvr1*) that received the same injury and were treated with either BLU-782 or ActA-mAb were collected at 5 dpi only. Muscle was then dissociated by mincing (3 and 5 dpi) or by using the "teasing method" (uninjured and 1 dpi; see Materials and Methods) and enzymatically digested for up to 1 hr. R206H-FAPs were isolated by FACS as live CD31-CD45-PDGFRα+SCA-1+GFP+tdTomato- cells, which selected for Tie2-lineage-labeled FAPs (green) that were recombined at the *Acvr1*$^{tnR206H}$ locus (not red). FAPs were immediately lysed, total RNA isolated, mRNA captured, and cDNA libraries prepared using a strand identity-preserving method. Approximately 30 million 75 bp, single-end reads were generated per sample.

**Supplemental Figure 3**

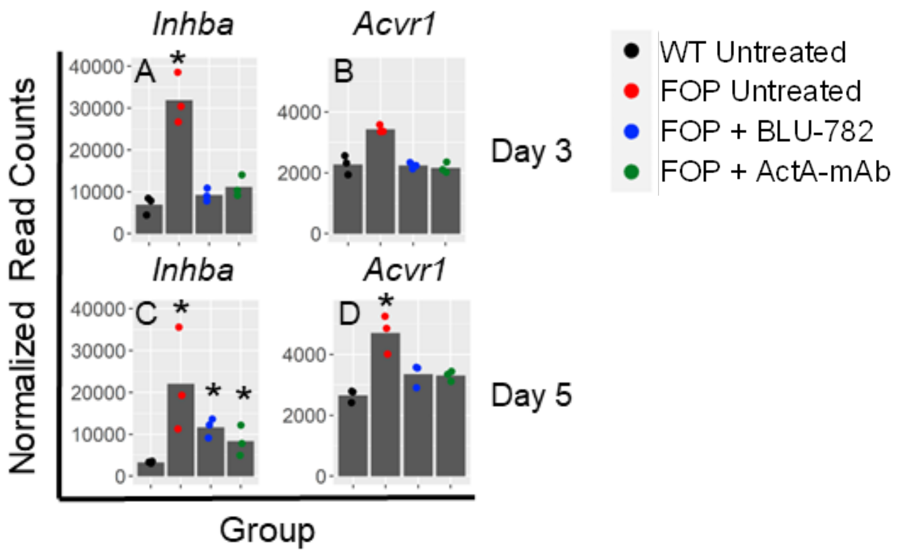


**Figure S3. *Inhba* and *Acvr1* are upregulated in R206H-FAPs after muscle injury.** R206H-FAPs isolated from injured muscle of untreated FOP mice exhibited > 4-fold upregulation of *Inhba* at both 3 dpi (A) and 5 dpi (C). *Acvr1* expression was significantly upregulated at 5 dpi (D) but not 3 dpi (B). Both *Inhba* and *Acvr1* expression levels were reduced in R206H-FAPs treated with either BLU-782 or ActA-mAb, although *Inhba* levels remained statistically elevated compared to controls at 5 dpi (C). Asterisks indicate a significant difference compared to controls, as determined by an adjusted p-value < 0.05 using a Wald test with Benjamini-Hochberg correction (DESeq2), and either 50% higher or 33% lower mean expression.

**Supplemental Figure 4**

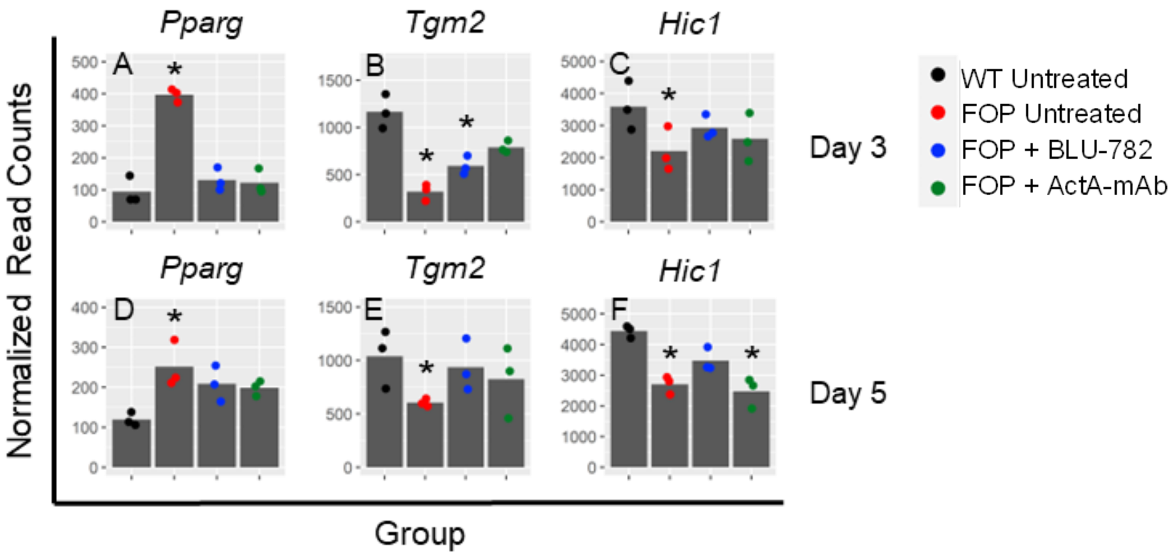


**Figure S4. R206H-FAPs upregulate a master adipogenic gene and downregulate retinoic acid targets after injury.** (A, D) The adipogenic transcription factor *Pparg* was upregulated at both 3 and 5 dpi in R206H-FAPs derived from untreated FOP mice. Treatment with each drug restored (A) or partially restored (D) control levels of *Pparg* transcripts. (B, C, E, F) Two retinoic acid targets, the apoptosis mediator tissue transglutaminase *Tgm2* (B, E), and the cell proliferation regulator hypermethylated in cancer-1, *Hic1* (C, F), were downregulated at both post-injury time points. Variable responses to drug treatments were observed, with a trend towards restoring control levels of *Tgm2* and *Hic1* in most cases. Asterisks indicate significance compared to controls, as determined by an adjusted p-value $< 0.05$ using a Wald test with Benjamini-Hochberg correction (DESeq2), and either 50% higher or 33% lower mean expression.
